# Rapid Changes in Chromosome Counts in Fishes from the Spiral Egg Clade within the Gourami Family (Osphronemidae)

**DOI:** 10.1101/2024.05.29.596523

**Authors:** Brendan Mobley, Andrew P. Anderson

## Abstract

Identifying clades with numerous and noticeable changes in chromosome counts is an important step in unraveling the evolutionary mechanisms that shape cytogenetic processes. Here, we describe low chromosome counts in a group of teleost fishes delimited by their unique spiral egg structure and with a species with a known low chromosome count within the labyrinthine clade (Osphronemidae). We sampled seven of nine known species within this spiral egg clade, reporting novel chromosome counts for five species and confirming two others. Overall, we find high variability in both chromosome count and arm number, which suggests a rapid loss of chromosomes during the emergence of the clade and numerous large-scale mutations occurring across evolutionary time. Lastly, we offer some possible explanations for these changes based on current and ongoing empirical and theoretical research. These data provide important information in cataloguing rapid chromosomal shifts in teleost fishes and highlights this group for further study in chromosomal and genomic evolution due to their karyotypic heterogeneity.

## Introduction

Variability of chromosome numbers across vertebrates and the evolutionary mechanisms that create it is an active point of inquiry for evolutionary biologists (Martinez et al. 2015). Finding clades with high variance or rapid changes presents valuable data points in resolving the various hypotheses on how karyotypes change over time. Some patterns can include changes due to long stretches of repeat content creating opportunities for mismatched recombination (Amores et al. 2014) such as found in mammals which range in chromosome counts from 2N=6 in the female muntjac deer (*Muntiacus muntjac*) (Wurster and Benirschke 1970; Graphodatsky et al. 2020) to 2N=102 in the plains viscacha rat (*Tympanoctomys barrerae*) (Gallardo et al. 2006; Stanyon and Graphodatsky 2012; Lebeda et al. 2020). Other patterns can involve microchromosomes as typically seen in birds which have a range from 2N=40 in *Ceratogymna bucinator* to 2N=136–142 in *Corythaixoides concolor* (Christidis 1990; Kretschmer et al. 2018). Conversely, non-avian reptiles are karyologically heterogeneous and exhibit distinct evolutionary trends between lineages (Deakin and Ezaz 2019) and have a narrower range of diploid chromosome counts than other groups (2N=24–70, Olmo 2005). Teleost fishes have been found to have notable karyotype evolution patterns, particularly with regard to extreme chromosome counts, rapid cytogenetic changes, or both (e.g. *Nothobranchius* 2N=16– 50:Krysanov et al. 2023 and *Corydoras* 2N=40–134: Shimabukuro-Dias et al. 2004) and have the widest 2N range of all vertebrates, ranging from 2N=12 in the marine species *Gonostoma bathyphilum* to 2N=446 in the freshwater species *Ptychobarbus dipogon* (Lebeda et al. 2020). Paradoxically, teleost fishes have a strong trend of conserved karyotypes (Galetti et al. 2000; Mank and Avise 2006; Nakatani et al. 2007) with over half of all karyotyped fish species having diploid chromosome counts of 48 or 50 (Mank and Avise 2006; Arai 2011), which has changed little from the proposed karyotype for the ancestor of all teleost fish (2N=52, Nakatani et al. 2007). With such diverse patterns across vertebrates, finding clades with unusual changes in chromosome counts is valuable to understanding chromosome evolution as whole.

An intriguing group of fishes with apparent chromosomal variance is within the family Osphronemidae, commonly called gouramis. The chocolate gourami, *Sphaerichthys osphromenoides*, has the lowest recorded chromosome count among freshwater fishes (Lehmann et al. 2021) with 2n=16 (<u>Calton and Denton 1974</u>). A species in the neighboring genus, the pikehead gourami, *Luciocephalus pulcher*, was reported to have 2n=20 (Arai 2011). Chromosome counts this low are exceedingly rare in fishes, as there are only thirteen fish species with a diploid chromosome count lower than 2N=22 (Lehmann et al. 2021). Furthermore, these counts are highly derived from the other Osphronemidae species, which generally have 2N values between 46 and 48 (Supplemental Table 1).

Both *S. osphromenoides* and *L. pulcher* are members of the “spiral egg” clade, a monophyletic group within the family Osphronemidae that includes the genera *Sphaerichthys, Luciocephalus, Parasphaerichthys*, and *Ctenops*. The monophyly was proposed based on the unique morphology of their eggs, which are covered in projections arranged in a spiral pattern, and later confirmed and refined with molecular evidence (Britz et al. 1995; Rüber et al. 2006). Another differentiating feature of the spiral egg clade is an angular jaw shape, which is taken to the extreme in the highly derived pike-like morphology of the piscivorous genus *Luciocephalus*. The spiral egg clade is also notable for having the only species, *S. osphromenoides* and *S. selatanensis*, in the family Osphronemidae with female broodcare via mouthbrooding compared to the overwhelmingly male mouthbrooders or bubble nesters in the family (Rüber et al. 2006), although recent evidence has called into question the sex of caring parent in *S. osphromenoides* (Zworykin et al. 2024). Chromosomes of the spiral egg clade remain largely uninvestigated; besides *S. osphromenoides* and *L. pulcher*, only one other species has been studied cytogenetically (*Ctenops nobilis*, 2N=44: Rishi et al. 1997). Given the low chromosome counts of *S. osphromenoides* and *L. pulcher* and the large 2N decrease relative to the wider family, we aim to characterize the karyotypes of additional members within the spiral egg clade. With this information we will describe the karyotypic diversity and evolutionary history for this remarkable group of fishes, thereby adding an extraordinary example to the chromosome count diversity in fishes specifically and animals in general.

## Methods

Fishes were sourced from the aquarium trade (Wet Spot Tropical Fish, Portland, Oregon, USA; Nationwide Aquatics, Tinley Park, Illinois, USA; Aqua Imports, Boulder, Colorado, USA), then held in species-specific tanks (110 liters) on a shared flow-through system (pH 7.0, GH 30 ppm, KH 40 ppm) with a 12/12 hour light/dark cycle with 30 minutes of dim light to simulate dawn and dusk. Specimens were housed for a minimum of one week before sampling to ensure good health for optimal cell proliferation.

Chromosome preparations were made following <u>Kligerman and Bloom (1977)</u> with the indicated modifications. Specimens were incubated in 0.005% colchicine solution for 6-7 hours, then euthanized and dissected to remove gill arches. Sex determination was conducted by gross examination of gonads with pictures taken throughout. Dissected specimens were stored in -80 °C for future molecular analyses. Gill arches were incubated in 0.4% KCl solution for 20-30 minutes, then fixed in two changes of 3:1 ethanol:acetic acid fixative for at least 30 minutes each, followed by an overnight fixation period at 4 °C. To prepare slides, tissue was homogenized into suspension by mincing in 50% acetic acid, then dropped onto a slide warmed to 30-40 °C and air dried. Slides were examined under phase contrast microscopy for quality control, then aged for at least one day at room temperature before being stained for 10 minutes in 10% Giemsa in pH 6.8 phosphate buffer (Gibco™ Gurr Buffer Tablets) and air-dried.

Chromosomes were examined under a Nikon Eclipse Ti-E microscope driven by Nikon NIS Elements AR software, then photographed with an oil immersion objective at 100x magnification and green color filtering using a Hamatsu ORCA-Flash4.0 camera. Digital images were optimized, then homologous chromosomes were paired by size and morphology and arranged in decreasing order of size using ImageJ v1.52v and Adobe Photoshop 24.3.0. At least 35 complete metaphase spreads were photographed from each specimen with completeness defined as the highest consistently observed chromosome count. Chromosomes were classified as metacentric (m), submetacentric (sm), subtelocentric (st), or acrocentric (a) according to their arm ratios (Levan et al. 1964). Chromosome arm number (Fundamental Number, FN) was calculated by considering metacentric and submetacentric chromosomes as biarmed and subtelocentric and acrocentric chromosomes as uniarmed.

## Results

We describe karyotypes for the first time in six species (Fig 1): *S. selatanensis, S. vaillanti, S. acrostoma, L. aura*, and *P. ocellatus*. We also confirmed the karyotypes of an additional two species (*S. osphromenoides, L. pulcher*) which matched those established in the literature (Calton and Denton 1974; Arai 2011). All species in the genus *Sphaerichthys* had different chromosome counts, with 2N ranging from 16–28 (Table 1). The number of chromosome arms (fundamental number, FN) showed less variation, with a range of 30–38. Notably, the sister species *S. osphromenoides* and *S. selatanensis* had a primarily biarmed and primarily uniarmed karyotype respectively, resulting in a nearly identical FN despite a 2N difference of 12. The other two sister species in the genus (*S. acrostoma* and *S. vaillanti*) had karyotypes that were a mix of biarmed and uniarmed chromosomes and had different values for both 2N and FN. We confirmed that *L. pulcher* had an entirely uniarmed karyotype of 20 acrocentric chromosomes. *Luciocephalus aura* had a primarily biarmed karyotype with six fewer chromosomes and eight more chromosome arms. *Parasphaerichthys ocellatus* had a primarily uniarmed karyotype with higher 2N and FN that were higher than any *Sphaerichthys* or *Luciocephalus* species, but lower than was reported for *C. nobilis* in the literature (Arai 2011).

**Figure 1.**
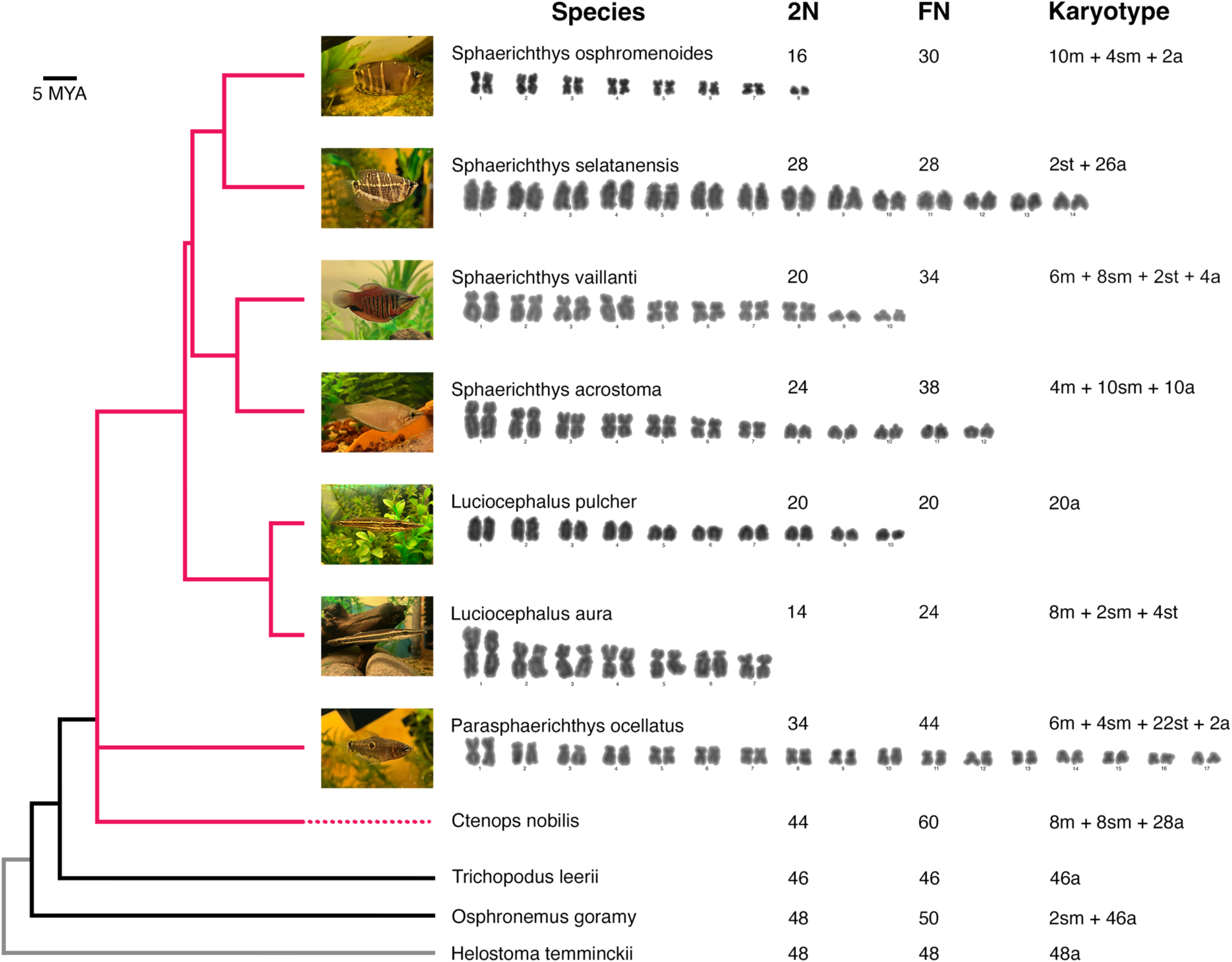
Selected anabantoid karyotypes. Phylogenetic relationships are from Ruber et al. (2006) and are shown to scale for the spiral egg clade (red) but not the selected species in the family Osphronemidae (black) or the outgroup (Helostomatidae, grey). Values for 2N, FN, and Karyotype for four species not generated in this study can be found in Arai (2011) and Grazyna et al. (2008). Karyotype formula describes number of metacentric (m), submetacentric (sm), subtelocentric (st), and acrocentric (a) chromosomes. Note that chromosome sizes vary widely between spreads and are thus not directly comparable except within the same spread. Live specimens were photographed with iPhone 13.

The two *S. selatanensis* that we sampled had different karyotypes (Supplemental Table 2). One had 28 uniarmed chromosomes (2N=28, karyotype 2st+26a), while the other had 26 uniarmed chromosomes and an unpaired metacentric chromosome (2N=27, karyotype 1m+2st+24a). The unpaired metacentric chromosome was approximately twice the size of the largest acrocentric chromosomes and may have been caused by a fused acrocentric pair. We cannot say whether this is a sex chromosome because we could not confidently determine the sex of either individual.

## Discussion

We found that the genera *Sphaerichthys, Luciocephalus*, and *Parasphaerichthys* have low chromosome counts (2N≤34) with high intra-genus variation in both 2N and FN. These trends are not observed in the karyotypes of the broader family Osphronemidae, which mostly have karyotypes similar to the hypothesized ancestral state for all teleost fishes (2N=52, Nakatani et al. 2007), thereby suggesting that rapid karyotype evolution occurred since the divergence of the spiral egg clade about 25 million years ago (Rüber et al. 2006). Karyotype evolution of this rate and magnitude has not been reported in teleost fishes. The drastic differences between karyotypes within the *Sphaerichthys* and *Luciocephalus* genera suggests that karyotype evolution may have played a role in speciation process by creating post-zygotic isolation (Canitz et al. 2016; Jackson et al. 2016; Mezzasalma et al. 2017; Romanenko et al. 2018), but it is also possible that the observed karyotype diversity happened alongside the speciation process instead of driving it (Krysanov et al. 2023). The observed karyotype pattern could have been created by the influence of genetic drift or other forms of neutral selection, extremely strong meiotic drive, the evolution of a trait that stimulates chromosome evolution, or a combination of these factors.

Genetic drift has been proposed to be the driving force behind the fixation of highly differentiated karyotypes in some clades of freshwater fishes, including the annual killifishes in the genera *Nothobranchius* (2N=16-50) (Krysanov et al. 2016, 2023; Krysanov and Demidova 2018) and *Aphyosemion* (2N=20-40) (Völker et al. 2005, 2007, 2008). These species tend to live in small, biogeographically isolated populations and frequently experience genetic bottlenecks and founder effects due to the ephemeral nature of their habitats (Völker et al. 2006; Krysanov and Demidova 2018; Krysanov et al. 2023), which may have sped up the accumulation of both intra-and inter-chromosomal mutations, with centric fusions being responsible for most decreases in chromosome count (Völker et al. 2005, 2008; Krysanov et al. 2016). *Sphaerichthys* and *Luciocephalus* species have very limited geographical ranges and, given that most of them are threatened or endangered according to the IUCN Red List, these populations are likely small; however, it is difficult to precisely estimate the strength of genetic drift on the evolution of the karyotypes in our study group because there is little information about their distribution and population structure.

An alternative explanation to genetic drift is meiotic drive. Species evolving under the influence of strong meiotic drive will tend to reach karyotypes of predominantly biarmed or uniarmed chromosomes (de Villena and Sapienza 2001; Molina et al. 2014). This has been observed in fishes and mammals, with the note that groups with high rates of mismatched karyotypes tended to have higher rates of chromosome evolution (Blackmon et al. 2019). The decrease in chromosome number in our study group relative to the rest of the family may have been caused by a meiotic drive toward biarmed chromosomes that rapidly fixed fusion mutations.

Additionally, the near-complete inversions in biarmed proportion between sister species (*S. osphromenoides* and *S. selatanensis, L. aura* and *L. pulcher*) are consistent with an inversion in the directionality of meiotic drive after or during the divergence of these species from their common ancestor, such that either fusions or fissions were preferentially fixed in one species but not the other. The karyotypes of *S. vaillanti* and *acrostoma*, which have a mix of biarmed and uniarmed chromosomes, may be partway through a shift to a completely biarmed or uniarmed karyotype. Additionally, changes in the arm number may have been caused by pericentric inversions, which may also be subject to the force of meiotic drive (Molina et al. 2014). The large differences in karyotype between recently diverged species indicates that the karyotypes were fixed extremely quickly, suggesting that meiotic drive would have to have been extremely strong if it was driving these changes. There are several counterbalancing forces that we would expect to weaken the strength of meiotic drive, including the relatively stronger force of genetic drift, as well as the general trend of changes in chromosome count tending to be slightly deleterious (King 1995).

Chromosomal rearrangements can have phenotypic impacts, particularly inversions, which can suppress recombination by capturing multiple locally adapted alleles (Kirkpatrick and Barton 2006; Berg et al. 2016; da Silva et al. 2021). Additionally, low chromosome number has been found to have correlations with phenotypic effects related to genome size (Gold 1979), including specialization (defined as being highly phenotypically derived from their close evolutionary relatives), tightening linkage groups, and occupying a narrower ecological niche (Gold 1979; Hardie and Hebert 2004). The observed rearrangements and reductions in the spiral egg clade may have played a role in acquiring highly specialized adaptations such as the ability of *Sphaerichtys and Luciocephalus* to live in peat swamp forests and the associated blackwater habitats, which are oligotrophic, sparsely inhabited, and highly acidic (pH < 4) (Polgar and Jaafar 2018). By contrast, *P. ocellatus* and *C. nobilis* are not adapted to such harsh conditions and are typically found in small muddy streams and pools. Additionally, it is possible that the rapid genomic rearrangements observed in this group may have contributed to the observed phenotypic differences in this subfamily, such as the highly derived morphology in *Luciocephalus*.

The spiral egg clade presents an excellent opportunity to understand how these exceptionally differentiated karyotypes arose and could give insight into larger patterns of chromosomal evolution. Advanced cytogenetic techniques could help clarify which types of chromosomal rearrangements occurred (ex. Ag-NOR staining, c-banding, FISH, etc.) as has been done in other species in the family Osphronemidae (<u>Grazyna et al. 2008; Pazza et al. 2009; Chaiyasan et al. 2021; Supiwong et al. 2021</u>), and measuring genome size of our study species would allow testing for non-conservative mechanisms of chromosome evolution. To test the influence of meiotic drive, work could be done to examine kinetochore protein levels during meiosis (Chmátal et al. 2014), as well as the amount of minor satellite DNA repeats on the centromere, which have been associated with the action of meiotic drive in the western house mouse (Iwata-Otsubo et al. 2017; Dudka and Lampson 2022). Other factors could be investigated that are known to stimulate chromosomal rearrangements such as repetitive DNA content (King 1995; Martinez et al. 2017) which was also implicated in the high incidence of chromosomal mutations in *Nothobranchius* (Krysanov et al. 2023). There are other monophyletic clades in the family Osphronemidae that have unusually differentiated chromosomes (Srisamoot et al. 2021), suggesting that the underlying mechanism driving karyotypic change in the spiral egg clade may be a shared ancestral trait and allowing for comparative genomic studies between the spiral egg clade and closely related groups. Finally, it is also worth noting that most Osphronemidae species have not been examined cytogenetically, hence karyotyping more species in the family Osphronemidae could reveal more clades with high karyotype differentiation. Further attention should be pair to this cytogenetically diverse group, as they could help resolve outstanding evolutionary questions of chromosomal rearrangements and diversity.

## Supporting information

Review of gourami karyotypes

Summary of karyotype observations in this study

## AUTHOR CONTRIBUTIONS

**Mobley**: Conceptualization; Investigation; Methodology, Data analysis and curation; Writing – original draft. **Anderson**: Conceptualization; Methodology; Data analysis and curation; Writing – review and editing; Supervision.

## ACKNOWLEDGEMENTS

This work was supported by National Science Foundation Grant #2147567. AP Anderson is supported by the NSF Postdoctoral Research Fellowship in Biology Grant #2010841. B Mobley was supported by a REP supplement to NSF grant #2147567.

## DATA AVAILABILITY

Raw images of chromosome spreads and finalized karyotype images can be found here: https://github.com/AndersonDrew/Gourami_Chromosome Contact authors for any additional data/information.

